# Methodological inconsistencies hinder comparative studies of alliance in the genus *Tursiops*

**DOI:** 10.64898/2026.01.14.699404

**Authors:** Hibiki Nishitani, Tadamichi Morisaka

**Affiliations:** Graduate School of Bioresources, Mie University, Mie, Japan

**Keywords:** alliance, cooperation, mammal, social behavior, toothed whale, *Tursiops*

## Abstract

Males of the genus *Tursiops* generally form alliances to cooperate for access to females, with variations in formation and complexity across regions. Given the variation in alliances, it may be possible to identify the factors that favor alliance formation through a cross-population comparative analysis. However, the association criteria that determine alliance formation in these previous studies are inconsistent, which has prevented researchers from deciding whether the variation in alliances reflects the inherent characteristics of each population or is simply pseudo-variation caused by criteria differences. To ensure the validity of future comparative analyses, we investigated the extent to which inconsistencies in association criteria among studies affect the variation through a literature review. We reviewed 50 studies from 23 regions and quantified criteria inconsistencies. We found discrepancies between the alliance statuses reported in the original studies and the outcomes of analyses in which criteria were exchanged within each study. Our results suggest that although inconsistencies in association criteria currently have little influence, the possibility that inconsistencies in criteria account for pseudo-variations in the alliance formation cannot be excluded. Based on this suggestion, we recommend that alliances should be determined by criteria that integrate behavioral data in addition to association data.

## 1 Introduction

Generally, research on coalitions and alliances in mammals follow the definition proposed by Harcourt and de Waal (1992), which defined cooperation as “an acting together of two or more individuals such that at least one amongst them stands to gain benefits unavailable thorough solitary action,” coalition as “a one-time cooperative action by at least two individuals or units against at least one other individual or unit,” and alliance as “long-term cooperative relationship, that is, partnerships that form coalitions on a regular basis.” Currently, coalition and alliance have been documented in mammals of six orders (Primates, Artiodactyla, Perissodactyla, Proboscidea, Carnivora, and Cetacea) and over 60 species (e.g., Sterck et al. 1997, Olson and Blumstein 2009, Smith et al. 2010, Bissonnette et al. 2014, Bissonnette et al. 2015, Lukas and Clutton-Brock 2018, Smith et al. 2022, Smith et al. 2023). Given that coalitions and alliances are regarded as one of the most complex social traits, cross-species comparisons of these traits in mammals are crucial for understanding the evolution of social behavior and cognitive abilities from both phylogenetic and adaptive perspectives (van Schaik 1989, Sterck et al. 1997, Connor 2007).

The males of two species of the genus *Tursiops* (bottlenose dolphin: *T. truncatus* and Indo-Pacific bottlenose dolphin: *T. aduncus*) generally form alliances to access females, with variations in formation (forming vs. non-forming) and complexity (uni-level or multi-level) across populations (Connor et al. 1992a,b, Möller et al. 2001, Wells 2014, Connor and Krützen 2015, Ermak et al. 2017, Connor et al. 2022, Brightwell and Gibson 2023). The society of *Tursiops* is characterized by a feature called fission-fusion dynamics, where spatial cohesion, group size, and composition change temporally (Connor et al. 2000, Auriel et al. 2008). Within this fluid society, males form distinguishable social relationships known as alliances. Allied males cooperate in the context of reproduction, such as mating with females, and defending or thieving females from other males or alliances (Connor et al. 2000). Indeed, genetic analyses has shown that alliance formation enhances reproductive success at the individual and/or alliance levels (Krützen et al. 2004, Wiszniewski et al. 2012b, Gerber et al. 2022, Duffield and Wells 2023). Thus, selection seems to favor alliance formation among males. Nevertheless, Brightwell and Gibson (2023) synthesized findings on *Tursiops* alliances across 31 regions and showed remarkable regional variation in both the formation and complexity of alliances. For instance, *T. aduncus* in Shark Bay, Australia (Connor et al. 1992a,b) and *T. truncatus* in Sarasota Bay, USA (Owen et al. 2002) form alliances, whereas *T. truncatus* in the Shannon Estuary, Ireland, does not (Barker et al. 2020). Additionally, in Sarasota Bay, alliances typically are formed as uni-level consisting of two individuals (Wells, 2014), whereas in Shark Bay and *T. truncatus* in the St. Johns River, USA, alliances are formed as multi-level consisting of four or more individuals or multiple alliances (Connor and Krützen, 2015, Ermak et al. 2017, Brightwell et al. 2025). Given the variation in alliances in *Tursiops*, it may be possible to identify the factors that favor the formation and complexity of alliances through cross-population comparative analysis (Connor et al. 2000, Whitehead and Connor 2005, Möller 2012).

However, the criteria that determine the formation and complexity of alliances in these previous studies are inconsistent, which has prevented researchers from deciding whether the variation in alliances reflects the inherent characteristics of each population or is simply pseudo-variation caused by criteria differences. Brightwell and Gibson (2023) showed that the criteria applied to determine the formation and complexity of alliances vary among previous studies. They pointed out that inconsistencies in the criteria applied among regions are likely to cause variations in alliances. However, because their research primarily aimed to organize current knowledge and test promising hypotheses about the alliance of *Tursiops* (Whitehead and Connor 2005, Möller 2012), the influence of inconsistencies in criteria was not evaluated in detail. If the variation in alliances is largely caused by inconsistencies in the criteria applied by each study, it is difficult to interpret the results of the comparative analysis from an ecological and evolutionary perspective. Therefore, before conducting a comparative analysis, it is necessary to evaluate the extent to which inconsistencies in the criteria applied among studies cause variations in the formation and complexity of alliances in *Tursiops*.

In previous studies, the concept known as “association” has played a central role in determining alliance formation. This concept assumes that individuals found together in a group will interact (Samuels and Tyack 2000, Whitehead 2008). Given the difficulty of directly observing social interactions in free-ranging cetaceans, association has commonly been used as a proxy measure. When association is applied to alliance research, it is assumed that long-lasting associations are correlated with functional behaviors that enhance reproductive opportunities, and that individuals who frequently associate are inferred to form alliances. However, two concerns arise from this inference. The first concern is the validity of the assumption that stable association serves as a proxy for functional behavior. Indeed, some studies in mammals suggest that this assumption does not hold universally (e.g., Connor and Mann 2006, Connor and Krützen 2015, Kawazoe 2021, Weiss et al. 2021, Fox et al. 2023). The second concern relates to the definition of “stable association”. According to Brightwell and Gibson (2023), the definition of stable association varies among researchers, but the consequences of such variability have not been evaluated. Therefore, evaluating the validity of these two lines is essential for conducting robust comparative analyses.

In this study, to ensure the validity of future comparative analyses of alliances in *Tursiops*, we investigated the extent to which inconsistencies in criteria among studies affect variation specifically in alliance formation through a literature review. Initially, we quantified the degree of inconsistency in the methodologies and criteria for alliance formation across previous studies. Subsequently, we evaluated the validity of the assumption that stable association serves as a proxy for functional behavior. Finally, we evaluated the reproducibility of alliance status under varying conditions of association criteria application.

## 2 Methods

The reporting and writing of review contents followed PRISMA -EcoEvo guideline (O’Dea et al. 2021) as a reference. However, as data collection had been completed before the guideline was consulted, the registration and study selection processes were not fully rigorous.

### 2.1 Classification and terminology

Following, Committee on Taxonomy (2023), we considered *T. truncatus* and *T. aduncus* as species, whereas all remaining species in the genus *Tursiops* were described as *Tursiops* sp. Although the taxonomy of the genus *Tursiops* varies among sources (Wilson and Reeder 2005, Mammal Diversity Database 2023), all sources list *T. truncatus* and *T. aduncus* as species.

In this study, we use only the term “alliance” in accordance with established convention by researchers in Shark Bay. Research on alliances has been spearheaded by studies in Shark Bay, and most researchers appear to use the terminology established in that region as a baseline. According to Harcourt and de Waal (1992), coalition and alliance are distinct concepts, but researchers in Shark Bay do not differentiate between these terms, using only the term alliance (see Connor et al. 2011, Connor et al. 2022).

In this study, we assumed that the primary benefit of alliance formation in *Tursiops* is to enhance the chances of mating opportunities (Connor et al. 2000). This assumption does not exclude other potential benefits such as improving predation avoidance and foraging efficiency (Connor et al. 2000). Following previous studies, we collectively referred to behaviors which related to alliances as “functional behavior”. It should be noted that, within the multi-level alliance structure conceptualized by researchers in Shark Bay (Connor and Krützen 2015), the functions of alliances differ according to level. Interactions between allied males and females are associated with lower-level (i.e. first-order alliances), whereas interactions between males or alliances, such as theft or defense over females, are primarily associated with higher-level alliances (i.e. second- and third-order alliances). Because this study focuses on alliance formation rather than complexity, we regard alliances as units that function in the context of reproduction and refer to any unit involved in enhancing mating opportunities as an alliance, regardless of its structural complexity.

### 2.2 Data collection

We collected the literature that exclusively focused on peer-reviewed articles and academic monograph. First, we added articles that we had already read to the database. Second, using Google Scholar and a subset of queries, we added relevant literature to the database. We used queries “Tursiops,” “alliance,” and “coalition” and added the literature displayed on page p.1 to p.40 on the Google Scholar to the database (search date is 2023/5/25). Third, we added literature in Table 1 of Brightwell and Gibson (2023) to the database. At the 25th Biennial Conference on the Biology of Marine Mammals held in 2024, we asked researchers about papers published this year and added those papers to the database. We then screened the content of the literature and its references in the database and extracted the subject literature that addressed alliance formation. These processes were conducted by the author NH.

**Table 1.**
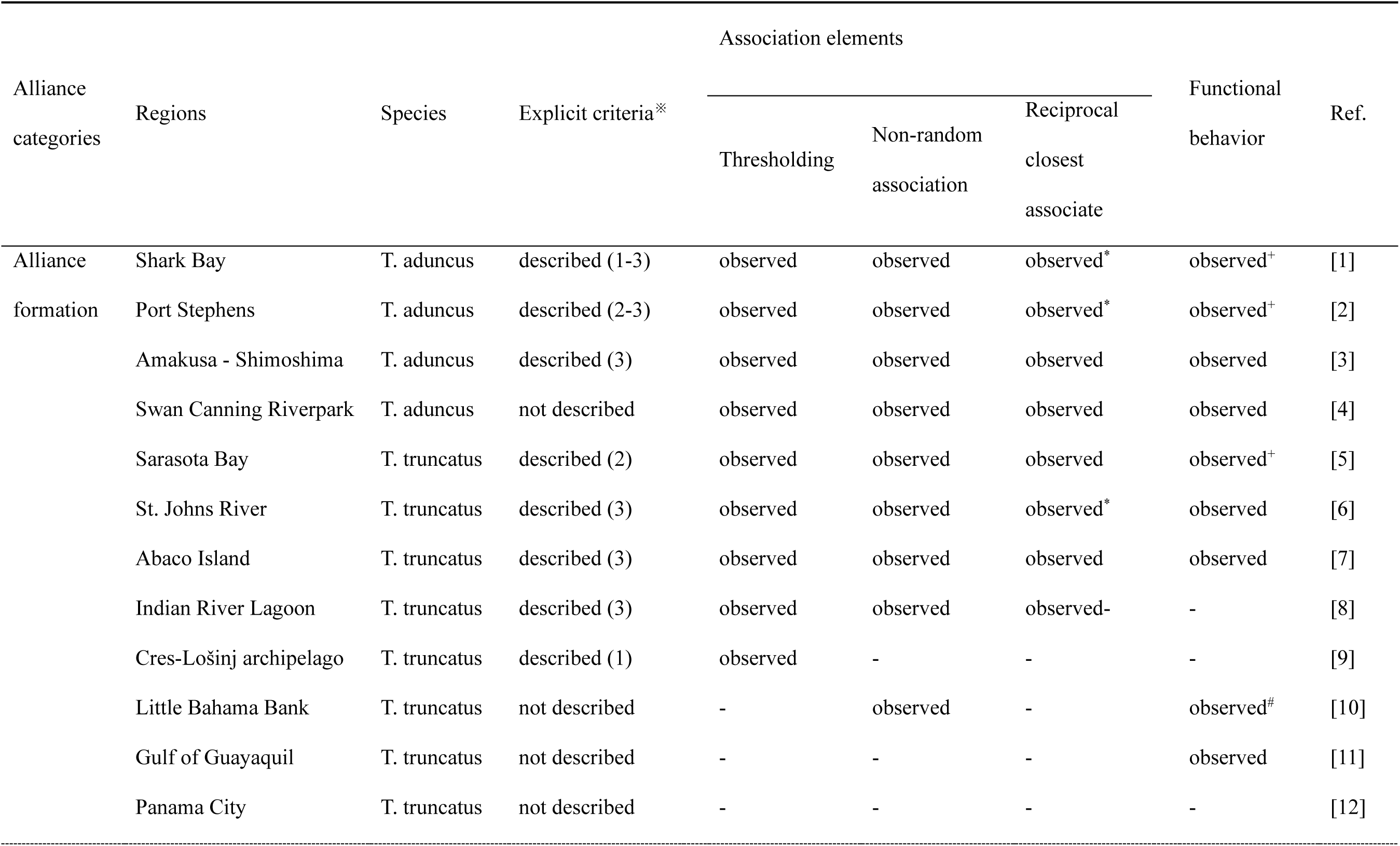
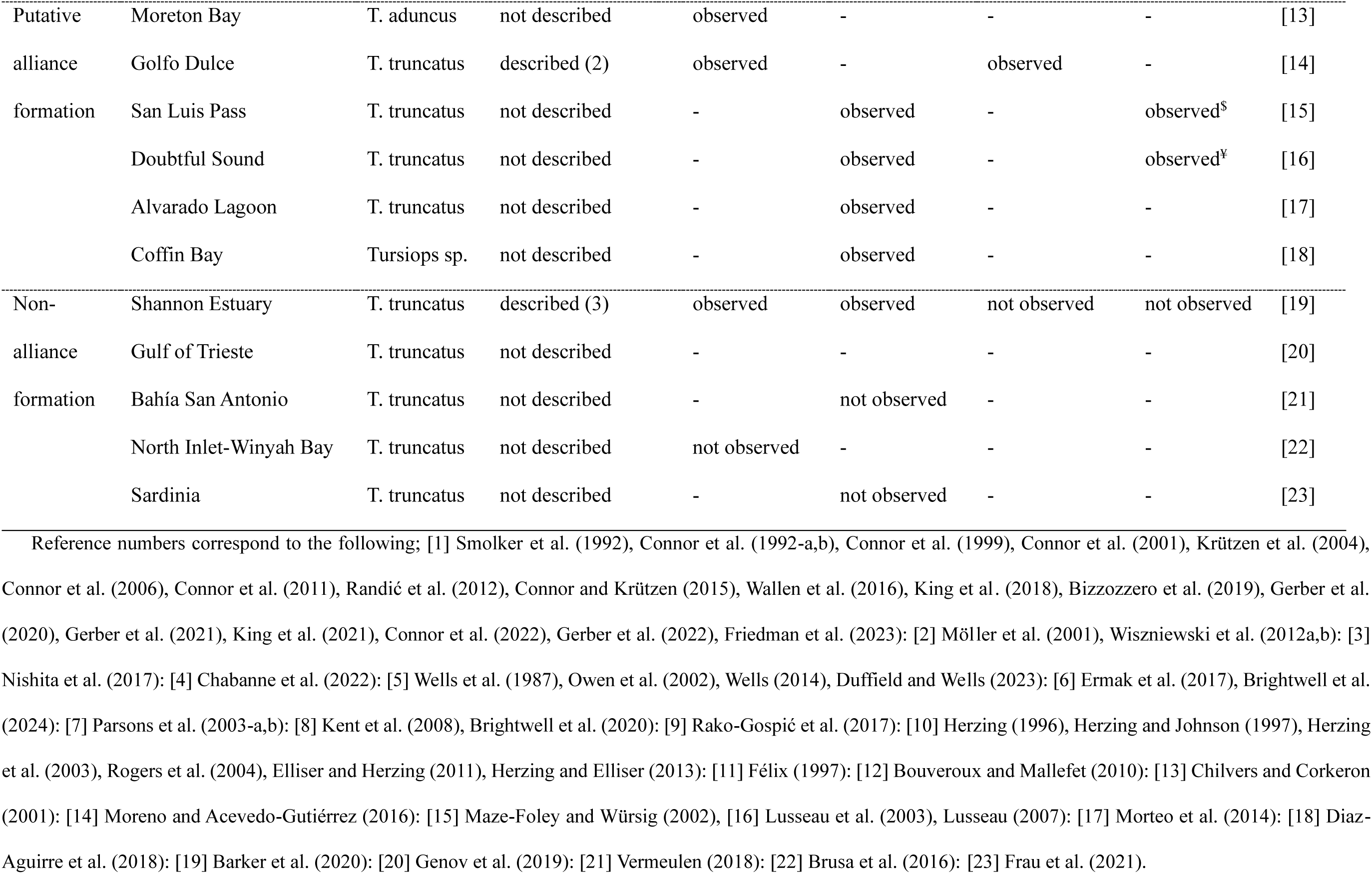
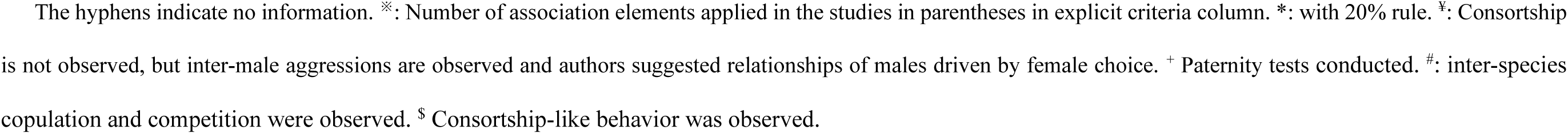
Alliance categories, species, explicit criteria, three association elements, and observation of functional behavior in the 23 regions.

### 2.3 Data process and analysis

First, we organized the status of the alliance formation in each region. Based on the content of each study, we classified the status of alliances in each region into the following three categories:

1. Alliance formation
  A. Alliance formation was declared in the study based on explicit criteria.
  B. Alliance formation was declared in the study without explicit criteria.
2. Putative alliance formation
  A. Alliance formation was suggested in the study with the terms “alliance-like,” “resemble alliance,” or “long-term relationship.”
  B. Alliance formation was suggested in the study, but the authors noted that additional evidence is required to confirm it.
3. Non-alliance formation
  A. Alliance formation was negated using the terms “absent” and/or “no evidence.”

The most recent literature was used to classify the regions into alliance categories if they had multiple sources.

Subsequently, we quantified the degree of inconsistency in the methodologies and criteria for alliance formation across previous studies. Referring to Brightwell and Gibson (2023), the criteria for determining alliance comprise two elements: a stable association (the two individuals be together for long time) and the observation of functional behavior. Thus, we focused on these two elements. Referring to previous studies (e.g., Connor et al. 1992, Möller et al. 2001, Owen et al. 2002, Ermak et al. 2017, Barker et al. 2020, Brightwell and Gibson 2023, Brightwell et al. 2025), we classified the criteria used to discriminate stable associations into three classes: 1) Thresholding: discriminating stable associations by the presence of males that showed a higher association index (the proportion of time that a dyad spends together) than a predetermined threshold. For instance, the threshold was set to 0.5, 0.8, or twice the population average. 2) Non-random association: discriminating stable associations by the presence of the males that showed a higher association index than expected by chance. 3) Reciprocal closest associates: discriminating stable associations based on the presence of males that showed the highest association index. A 20 percent rule that deals with a triplet, quartet, or more is sometimes applied when individuals do not have a reciprocal closest associate (see Smolker et al. 1992, Connor et al. 1992a,b, Möller et al. 2001). These three classes of association may be applied individually or in combination with one another. We recorded these three classes of associations as stable association elements. In addition, we recorded whether each region sets explicit criteria to determine the alliance formation.

We recorded the methods related to associations in each study, including the definitions of associations and groups and the quantification of associations (details in supplemental material). Since direct recording of interactions is difficult in free-ranging cetaceans, the “gambit of the group”, an assumption whereby all individuals observed within the same group are assumed to be associated, is widely applied. Under this assumption, the interpretation of association critically depends on how a group is defined. In the present study, when referring to cetacean research, we use the term “group” exclusively to denote a sampling unit, rather than a social unit (Syme et al. 2022).

Given that reports of functional behaviors are predominantly reported in Shark Bay, we recorded the types of functional behaviors described among frequently associating males in each region. Based on previous studies, we assumed that alliances enhance mating opportunities through tactics such as the maintenance of consortships mediated by allied males and direct inter-male or inter-alliance competition over females (Connor et al. 1992a,b, Connor et al. 1996, Scott et al. 2005, Connor and Krützen 2015). Following this, we focused on recording interactions from males to female(s) and interactions among males. It is noted that allied males have been reported to have larger home ranges than non-allied males (Owen et al. 2002; Wells 2014), suggesting that alliance formation may be functional through increasing encounters with females. However, Rako-Gospić et al. (2017) reported that allied males have smaller home ranges than non-allied males. Given these contradictory findings, we did not treat home-range expansion associated with alliance formation as a functional behavior. In addition, we did not treat sexual displays as functional behaviors, as they are more likely to function in strengthening social bonds among males (Hill-Cousins et al. 2025), rather than directly contributing to coerced consortships or inter-male competition.

To evaluate the extent to which inconsistency in criteria affect variation in alliance formation, we conducted two types of criteria-change analysis. The first analysis evaluated the validity of the assumption that stable association serves as a proxy for functional behavior. This analysis included only regions where behavioral data were available. We listed all possible association criteria that could be derived by combining three classes of association, resulting in seven distinct criteria. In the analysis, all seven association criteria were applied to each region, and the reassessment outcomes under each criterion were recorded. The results of analyses were then compared with the behavioral observations originally reported for each region. When the results and the observations of functional behavior agreed, either when the criterion was satisfied and the behavior was observed, or when the criterion was not satisfied and the behavior was not observed, the outcome was recorded as “positive”. When the results and observations disagreed, the outcome was recorded as “negative”. If the relevant association criteria had not been investigated in the original study and a determination could not be made, the outcome was recorded as “unclear”. We recorded the number of regions classified as positive, negative, or unclear under specific criteria.

The second analysis evaluated the reproducibility of outcomes under varying conditions of association criteria. In the analysis, seven possible association criteria were applied, and the reassessment outcomes under each criterion were recorded for each region. The results were compared with the alliance status reported in the original studies. In this analysis, regions classified as putative alliance formation were regarded as equivalent to those classified as alliance formation. When the results and the original alliance status agreed, the outcome was recorded as positive. When the results and the original status disagreed, the outcome was recorded as negative. If the relevant association criteria had not been investigated in the original studies and a determination could not be made, the outcome was recorded as unclear. We recorded the number of positive, negative, and unclear cases in each region under each of the seven applied criteria.

## 3 Results

We extracted and reviewed 50 literature sources from 23 regions. The number of regions examined for *T. aduncus*, *T. truncatus*, and *Tursiops* sp. were 5, 17, and 1, respectively.

Table 1 shows the alliance categories, explicit criteria, association elements, and functional behavior in each of the 23 regions. Twelve regions were classified as alliance formation, six as putative alliance, and five regions as non-alliance formation. Ten regions set explicit criteria that discriminate against the alliance formation, whereas the remaining 13 regions did not. The number of association elements embedded to the explicit criteria varied among regions. The numbers of regions that examined three, two, one, and none of the association elements were nine, one, ten, and three, respectively. Twelve regions reported functional behavior, while the remaining 11 regions did not. In the regions categorized as alliance formation or putative alliance formation, males satisfied any of the three association elements, except for two regions without examination of any of the three elements. In the three regions were classified as non-alliance formations, males did not satisfy any of the investigated association criteria, except for one region without examination of any of the three elements. However, one region, the Shannon Estuary, is categorized as a non-alliance despite the presence of males that satisfy two of the three association elements. In the 12 regions where observations of functional behavior were reported, such behaviors were observed in regions classified as alliance formation or putative alliance formation, but not in the non-alliance formation region.

Table S1 shows the definitions of associations and groups as well as quantification of associations used in each study. While definitions of groups sometimes differed among studies conducted in the same regions, the totals reported in the descriptive statistics below do not necessarily amount to 23. Groups were defined based on proximity (e.g., 10 m-chain rule, 100 m radius rule) in six regions, based on behavioral state in two regions, and using a combination of proximity and behavioral state in 13 regions. In three regions, no explicit group definition was identified. In 18 regions, associations were recorded based on group membership, that is, following the gambit of the group. In one region, associations were defined based on inter-individual distance, while in four regions no explicit description of how associations were defined was provided. The application of association indices also varied among studies within the same regions. The half-weight index (HWI) was used in 22 regions, the simple ratio index (SRI) in two regions, and the generalized affiliation index (GAI) in one region.

Table S2 shows functional behaviors were observed among males that frequently associated in each region. Eight regions reported males-female interactions. Detailed and structurally described behaviors were reported only for Shark Bay. In other regions, reports were based on citations of the Shark Bay literature (Port Stephens and St. Johns River), descriptions of observed behavioral elements (Amakusa-Shimoshima and Swan Canning Riverpark), or anecdotal accounts (Sarasota Bay and San Luis Pass). In the Shannon Estuary, it was reported that typical alliance behaviors were not observed based on citation of the Shark Bay literature. In addition, six regions reported males-male(s) interactions. Detailed description of aggressive behaviors among males were reported in Shark Bay and Abaco Island and others reported functional descriptions that listed the behavioral elements observed.

Figure 1 shows the first analysis evaluating the validity of the assumption that association serves as a proxy for functional behavior. When the applied criteria were changed, the number of regions recorded as positive ranged from 7 to 10 (median = 8, n=12) and recorded as unclear ranged from 1 to 4 (median = 4, n=12). Negative outcomes were only recorded under three conditions such as “Thresholding”, “Non-random”, and “Thresholding + Non-random” in one region, the Shannon Estuary. Figures 2 shows second analysis evaluating the reproducibility of alliance formation under varying association criteria. The results were obtained by applying all seven association elements across all regions and comparing them with the alliance status originally reported for each region. The mean positive rate across regions was 0.45 (SD = 0.43). The mean negative rate was 0.02 (SD = 0.09), and the mean unclear rate was 0.53 (SD = 0.44). Negative outcomes were only recorded in three conditions such as “Thresholding”, “Non-random”, and “Thresholding + Non-random” in one region, the Shannon Estuary.

**Figure 1.**
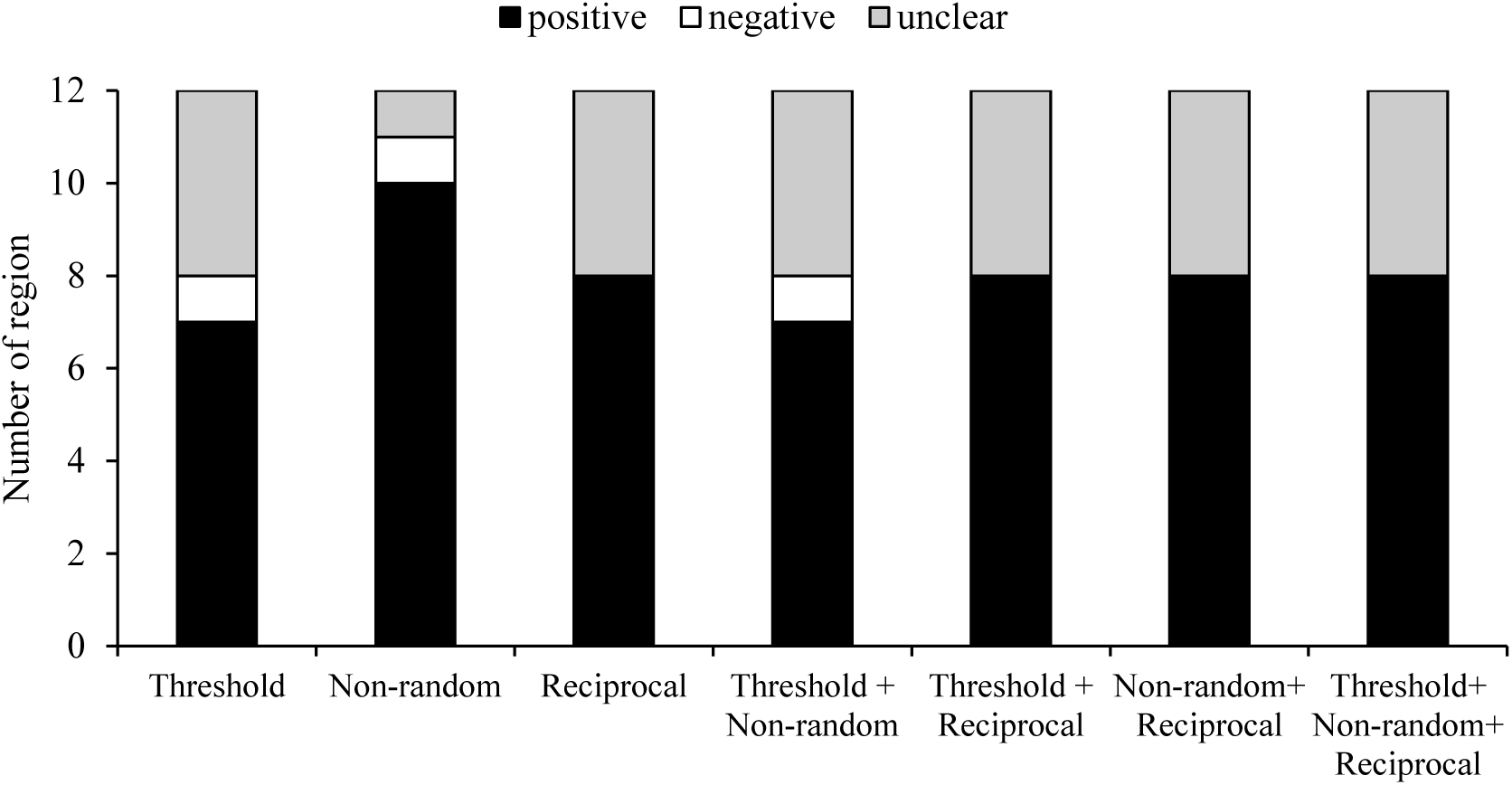
Results from the first analysis evaluating the validity of the assumption that association serves as a proxy for functional behavior. The number of regions classified as positive, negative, or unclear under specific applied criteria are shown.

**Figure 2.**
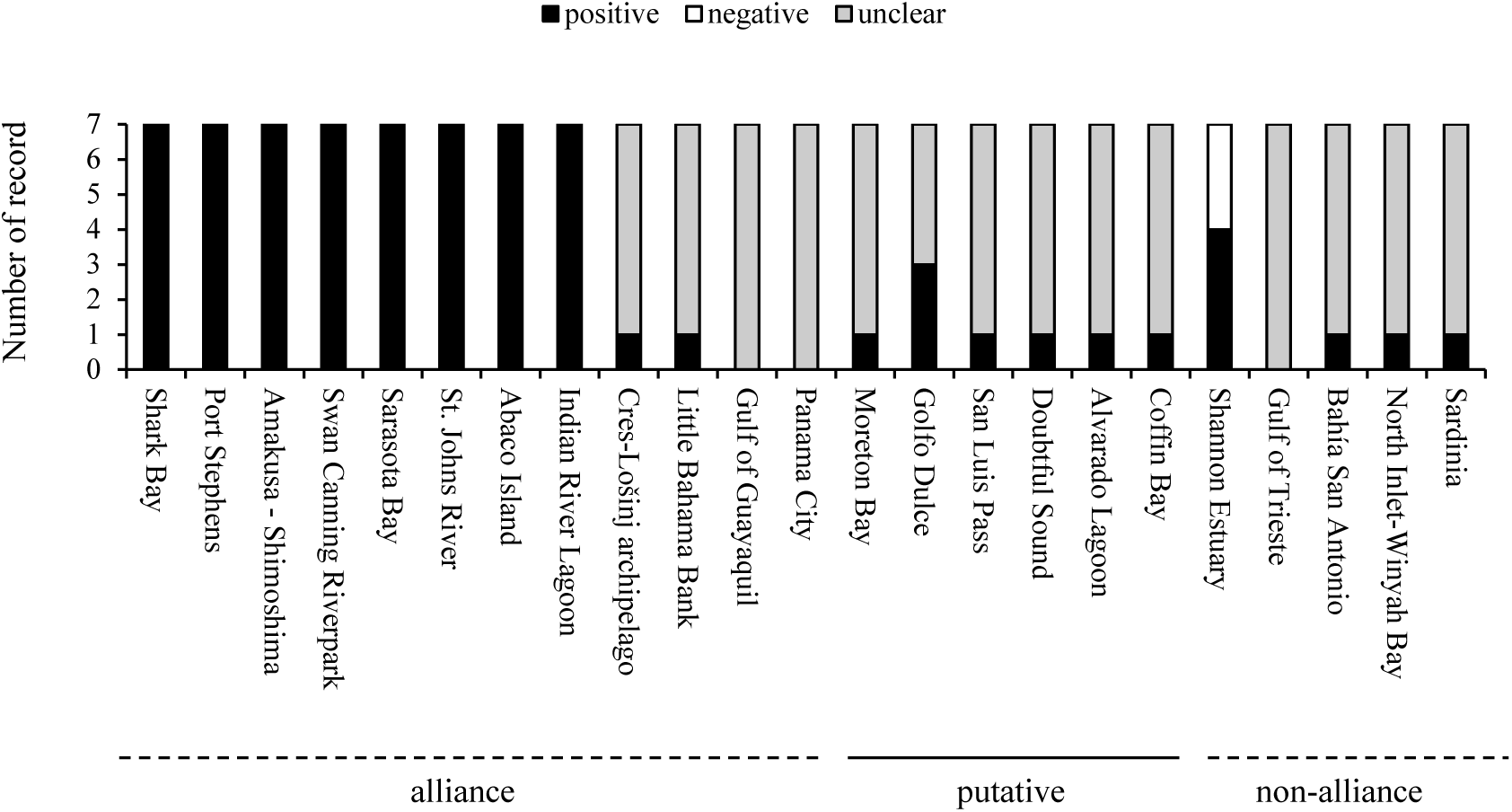
Results from the second analysis evaluating the reproducibility of alliance formation under varying association criteria. For each region, the outcomes obtained when seven different association criteria were applied were recorded as positive, negative, or unclear.

## 4 Discussion

Through a literature review on 23 regions, we quantified the inconsistencies in the methodologies and criteria used to determine alliance formation. The first analysis showed that, although association generally serves as a relatively good proxy for functional behavior, negative outcomes were observed in one region under specific criteria, where association criteria were satisfied but functional behavior was not observed. Given that the unclear rate reached approximately 30% and that functional behaviors have not yet been reported in about half of the regions, the possibility that the negative rate is underestimated cannot be ruled out. The second analysis showed that the reproducibility of alliance formation is generally unclear, and negative outcomes were observed under certain conditions. These findings suggest that variations in the applied association criteria may lead to differences in the resulting determination for alliance formation. Taking together, our findings indicate that criteria inconsistencies cannot be entirely ruled out as a source of pseudo-variation in alliance formation. Our results support the discussion of Brightwell and Gibson (2023) that inconsistencies in the criteria applied across regions are likely to lead to variation in alliance formation. Based on these findings, we discuss methods for determining alliances in a practical way that facilitates interpretation in future comparative research.

### 4.1 Difficulty to determine for alliance formation using association alone

Determining alliance formation using association criteria alone is likely challenging. First, although our results suggest that association is reasonably supported as a proxy for functional behavior, they also indicate that its validity depends on the criteria. Currently, this dependency is based solely on a single case, and both its extent and strength remain unclear. In line with the concerns raised in the present study, it has been questioned whether association can serve as a substitute for interaction and/or cooperation between individuals (Gowan 2019). For example, in Shark Bay, although alliances were defined based on associations and functional behavior during the 1990s, first-order alliances that vary in stability within second-order alliances are now defined solely based on functional behavior rather than association (Connor and Krützen 2015). Additionally, in killer whales (*Orcinus orca*), the frequency of social interactions was negatively correlated with age, but this correlation was not found based on the strength of association and age (Weiss et al. 2021). Furthermore, in Japanese macaques (*Macaca fuscata*), the frequency of coalitions does not correlate with the strength of the association (Kawazoe 2021). Coalition among female chimpanzees (*Pan troglodytes*) appears to depend on social tolerance rather than strong social bonds (Fox et al. 2023). However, some studies have shown that association measures can serve as reliable proxies for interactions and cooperative behavior (Atlantic spotted dolphin, *Stenella frontalis*, Danaher-Garucia et al. 2022; bonobos *Pan paniscus*, Tokuyama and Furuichi 2016). These instances indicate that the various association criteria do not necessarily serve as substitutes for interactions and/or cooperation or delineate functional unit.

Second, our results indicate that the strictness of association criteria is arbitrary, suggesting that ensuring reproducibility may be difficult. For instance, males who satisfied all three association criteria were considered to alliances in the Shannon Estuary, whereas males who satisfied only one or two association elements were considered alliances in some of the other regions. If the strict criteria set in the Shannon Estuary, for example, were applied to other regions, the alliance status would likely change from its original one. Indeed, the moderate unclear rate emphasizes these significances.

In contrast to terrestrial studies, relationships among *Tursiops* have been described as alliances both with and without obvious functions inferred from behavioral observations. Coalition and alliance in terrestrial mammals is indicated by observing social interaction and/or cooperation as well as an association (e.g., lion, *Panthera leo*, Chakrabarti and Jhala 2017: fallow deer, *Dama dama*, Jennings et al. 2011: stump-tailed macaque, *Macaca arctoides*, Toyoda et al. 2022: bonobo, Tokuyama and Furuichi 2016: chimpanzee, Gilby et al. 2013, Feldblum et al. 2021: Japanese macaque, Kutsukake and Hasegawa 2005: spotted hyena, *Crocuta crocuta*, Strauss and Holekamp 2019). The key challenges faced in the study of cetacean social behavior are the difficulty of frequently observing behaviors in the natural situation. As a result, the use of association as a proxy has become an unavoidable necessity in practical research contexts (Samuels and Tyack 2000, Whitehead 2008). However, given that over half of the regions reported functional behavior and that no discrepancies were found alliance statuses determined solely based on functional behavior, behavioral data should be incorporated into methods for determining alliance formation, in addition to associations, to ensure both unambiguous interpretation and reproducibility.

### 4.2 Essence for determining alliance

Considering the above discussion, we reconsider the currently common inference for determining alliances and identify the essence required for this process. While association criteria are needed for practical research in cetaceans, determining alliances based solely on association criteria are accompanied by functional uncertainty and issues of reproducibility (Figure 3). To solve these issues, we emphasize it is essential that, at the early phase of each study, behavioral data must report in some form to demonstrate that males determined to be allied inferred by association criteria are involved in interactions or cooperation. While the quality of such behavioral data in previous studies can vary substantially (Table S2), reported allied males are reasonably interpretable and functionally comparable. To secure reproducibility, we recommend that researchers should define convenient criteria such as associations that may serve as proxies for actual interactions and cooperation in combination with the results of behavioral data (Fig. 3). For instance, Connor et al. (1992a,b) analyzed both association and functional behavior, and found functional behavior was only observed among males with exhibited association indices of 0.8 or more. Thus, they defined males exhibiting association indices of 0.8 or more as alliances. At present in Shark Bay, second order alliances are reasonability distinguished and defined using cluster analyses (Connor and Krützen 2015, King et al. 2021, Connor et al. 2022). Möller et al. (2001) incorporated behavioral observations into the criteria for determining alliances directly, in addition to using association indices.

**Figure 3.**
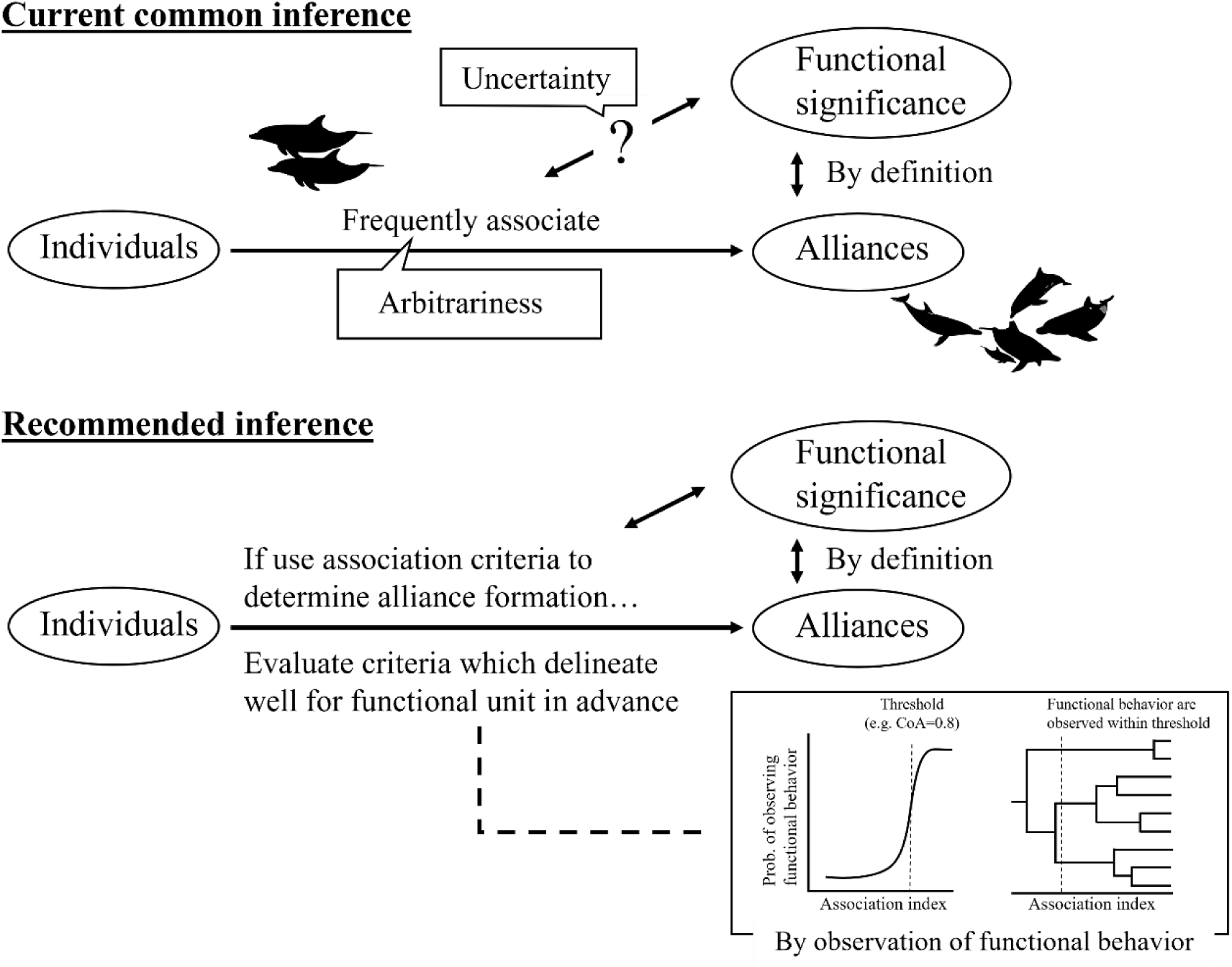
Conceptual scheme of currently common inference (above) and recommend inferences (below).

Standardizing specific methods is likely to be difficult, as they are strongly influenced by logistical constraints and the observational techniques available in each study. By allowing a degree of flexibility such as contents and amount of behavioral data and form of reporting, our recommendation enables each study to evaluate the validity of employing convenient association criteria, even when behavioral data are limited. It is essential that each study reports behavioral data with association data and defines criteria that can be reasonably interpreted as alliances by integrating these data. Accordingly, it contributes to resolving issues such as the ambiguity in interpreting alliances and the reproducibility of each study, with ensuring the feasibility of social research on cetaceans and future comparative analysis.

A potential criticism is that, in the case of *Tursiops*, there is currently no evidence that frequently associating males serve functions other than reproduction, and thus it may be reasonable to assume that such associating males gain some reproductive benefit. However, even if this criticism is correct, our results show that no universal association criterion has been established and applied to distinguish males who interact or cooperate. In other words, it addresses the functional uncertainty but not issues of reproducibility. To define the interpretable and reproducible definition based on association, behavioral observation is necessary.

### 4.3 Reconstruction of current findings

Following our recommendation, we consider that only social relationships thought to enhance mating opportunities, as supported by criteria with functional behavior, should deserve alliances. Given that males are generally considered less likely social than females in mammals (Clutton-Brock 2016), inter-male social relationships inferred solely from associations should nonetheless be regarded as valuable. Considering this, we recommend the term “social bond” to specifically refer to relationships based solely on association, distinguishing them from “alliances” which have exact functional meaning (Cords and Thompson 2017).

Referring to our results and recommendations, we reassessed previous studies as shown in Figure 4. We designated regions where functional behavior was observed as “alliance formation” (n=8). Additionally, we re-designated Shark Bay and St. Johns River as “multi-level alliances” (n=2, Connor et al. 2022, Ermak et al. 2017). Subsequently, we designated regions where functional behavior was not observed as “non-alliances” (n=1). We designated the regions where males who satisfy all three association criteria are observed or putative functional behavior are observed as “social bond” (n=4). Finally, we designated the remaining regions as “unknown” (n=10). Note that we arbitrarily applied the strictest criteria to determine the “social bond” Thus, if the criteria were revised, the status of social bonds in each region would vary.

**Figure 4.**
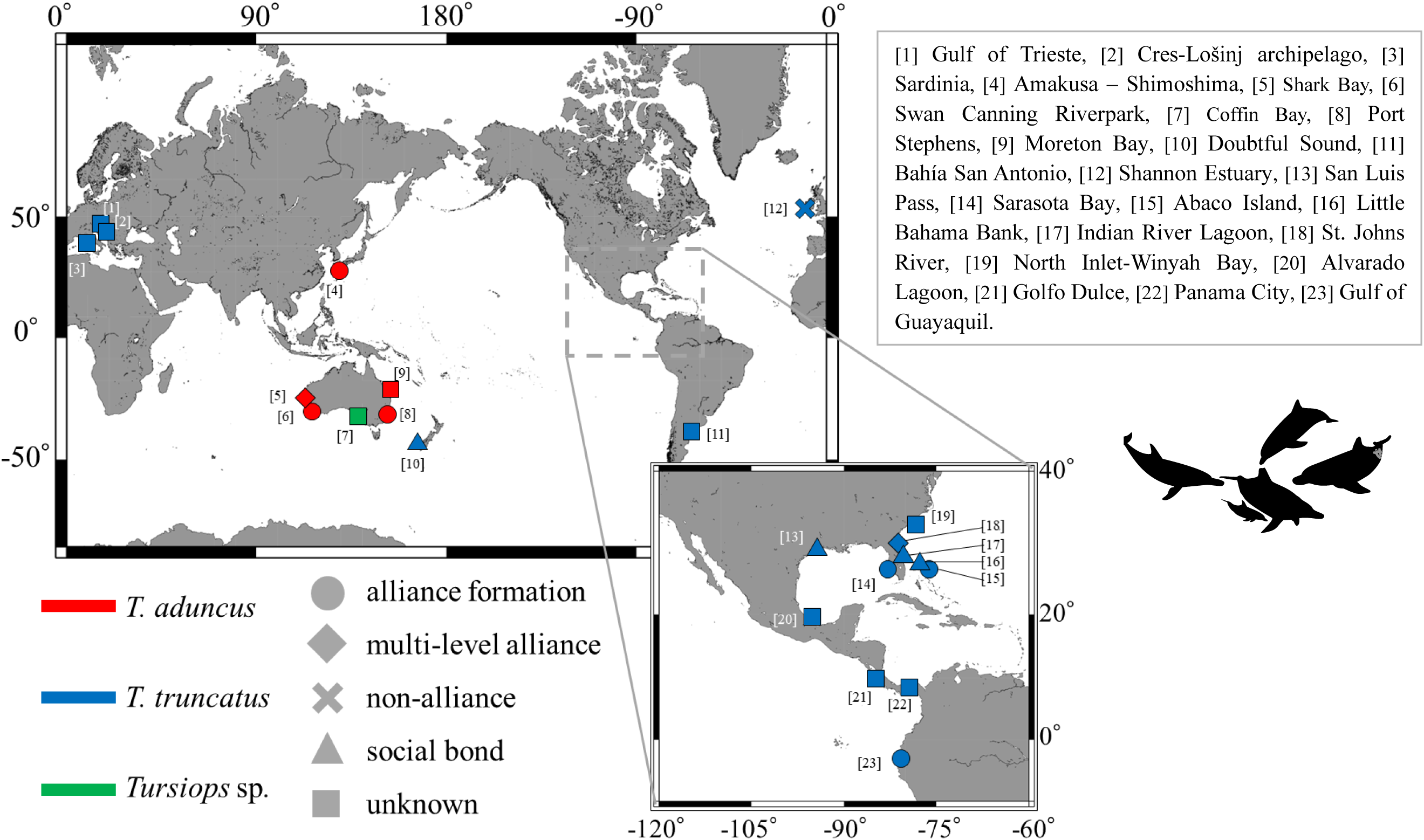
Reassessment of previous studies on the formation and complexity of alliances in *Tursiops*.

### 4.4 Cautions and limitations

There are several considerations regarding determining alliances. First, the likelihood of observing functional behavior depends on both the field method and frequency of field surveys. In cetacean research, two main approaches are commonly employed. “Survey”, which allows for the coverage of a broad area and the detection of multiple groups but provides limited observation time per group; and “focal follow”, which enables continuous tracking of a specific group, but it is challenging for researchers to allow for the observation of multiple groups (Mann 1999). If the former method is employed at low frequency, there is a risk of false negatives for recording behavior. Referring to the methods of studies classified as alliance formation in Figure 4, we recommend that data with a minimum of 100 survey days or 200 sightings, and at least five identifications per dyad, be used for determining alliance formation in order to reduce the likelihood of such errors.

Next, we turn to the analysis of association. Although it is common practice in cetacean research to define association based on co-identification within a group, a longstanding issue is how to define a “group” as a sampling unit (Syme et al. 2022). While definitions of the group in each study listed on Table S1, we were unable to evaluate the extent to which differences in group definitions may have influenced variation in alliance formation. Given that the active space, the spatial scale at which dolphins perceive group membership, is expected to vary according to environmental conditions (Janik 2000, Quintana-Rizzo et al. 2006, Sørensen et al. 2024), it is difficult to establish a universal group definition for alliance research. In addition, even if a theoretically suitable group definition is established, it may not be practically applicable in the field due to limitations in the observational scope of researchers (Syme et al. 2022). So far, we consider that the use of group definitions from previous studies is currently the most practical approach in comparative contexts. In addition, differences in the methods used to quantify association frequency may influence variation in alliance formation. Although most studies applied half-weight index (Table S1), we did not test the differences of quantifying the association that may affect the qualitative comparisons of alliances. The uncertainty of association frequency can be influenced by multiple factors, ranging from sampling issues such as missed individual identifications to demographic processes such as immigration, emigration, and mortality. Accordingly, applying the most appropriate quantification methods on a case-by-case basis, tailored to their specific survey protocols and the demographic characteristics of their study populations (see Cairns and Schwager 1987, Croft et al. 2008, Whitehead 2008, Farine and Whitehead 2015, Whitehead and James 2015, Hoppitt and Farine 2018).

### 4.5 For future comparative studies

In this study, we assumed that the maintenance of consortship constitutes functional behavior, but this assumption was primarily confined to cases where it is maintained by males. However, the possibility that consortships may also be maintained through mechanisms initiated by females, such as mate choice, remains hypothetical and open to further studies (Connor et al. 1996, Connor et al. 2000, Lusseau 2007, Wells 2014. see Table S2). If there are multiple pathways by which consortships are maintained, the definition of functional behavior adopted in this study may not have been comprehensive. A deeper understanding of the function of male alliances is required for establishing a more appropriate definition. To achieve this, it is necessary to observe behavior with association and this consideration does not conflict with our recommendation. In future research, the integration of standard direct behavioral observations, whether from boat-based or underwater, with complementary techniques such as unmanned aerial vehicle footage or hydrophone recordings is likely to be particularly effective (King et al. 2021, Chereskin et al. 2022, King and Jensen 2023, Chereskin et al. 2024).

In the present study, we did not examine the complexity of alliances, such as multi-level structure, size and stability. Nevertheless, the recommendation here may also contribute to enhance the study and comparison of alliance complexity. The conceptual framework for alliance complexity has been established exclusively through research in Shark Bay (Connor et al. 1992a, b, Connor et al. 1999, Connor et al. 2011, Connor and Krützen 2015, Connor et al. 2022). In summary, alliances in Shark Bay are conceptualized as comprising three-levels. Second-order alliances, typically formed by 4 to 14 males, maintain long-term associations over several decades and are considered the core social units. Within these second-order alliances, 2 to 3 males form first-order alliances. In addition, several first- or second-order alliances form third-order alliances. First-order alliances primarily function in maintaining consortships, while second- and third-order alliances play roles in inter-male conflict. Within this multi-level structure, there are two types of continuous variation. The size of second-order alliances ranges from 4 to 14 males, and the stability of first-order alliances within them varies along a continuum from labile to stable. Therefore, alliance complexity in Shark Bay is characterized by three elements: discrete levels, functional differentiation between levels, and continuous variation within levels.

To date, in regions where alliance formation has been reported, few studies have explicitly assumed this conceptual structure (but see Ermak et al. 2017, Brightwell et al. 2020, Brightwell et al. 2025). As a result, the distribution of multi-level structures remains unclear, and it remains difficult to identify its causal factors (but within-population comparative analyses have been conducted; see Connor et al. 2017, Sørensen et al. 2024). To apply the Shark Bay framework, it is first necessary to define multi-level structure, which requires identifying functional distinctions between them. This consideration again highlights the importance of behavioral observation, for which our recommendation provides a useful foundation. Once each level is defined, continuous variation within each level, such as alliance size and stability, can also be assessed. Determining alliance size also requires behavioral data to interpret accurately. For example, in cases where four males are allied, it is difficult to determine whether they form a single first-order alliance or two first-order alliances without behavioral information (e.g., Möller et al. 2001). In future research, determining alliance complexity may require the use of model selection approaches. These could include fitting several candidate alliance models, such as uni-level and multi-level models, to empirical data and selecting the model that provides the best fit to the observed data.

Currently, it is likely difficult to test the hypotheses concerning the variation in alliance among *Tursiops* through cross-regional comparisons (Connor et al. 2000, Whitehead and Connor 2005, Möller 2012). One of the reasons is the paucity of reports on the absence of alliances (Figure 4). Additionally, most previous studies have investigated only alliance formation, without addressing the complexity and size of alliances. Both issues are likely to be mitigated through the accumulation of data generated by studies incorporating behavioral observations. In addition to *Tursiops*, social relationships among males have been reported in other toothed whales (e.g., pantropical spotted dolphins, *Stenella attenuata*, Pryor and Kang-Shallenberger 1991; spinner dolphin, *Stenella longirostris*, Johnson and Norris 1994; Atlantic spotted dolphin, Herzing 1996, Elliser and Herzing 2014; *Sousa sp.,* Allen et al. 2017; Risso’s dolphin, *Grampus griseus*, Hartman et al. 2008, Hartman et al. 2020, Hartman et al. 2023; Cuvier’s beaked whale, *Ziphius cavirostris*, Cioffi et al. 2021; sperm whale, *Physeter macrocephalus*, Kobayashi et al. 2020, Pacific White-sided dolphin, *Aethalodelphis (formarly Lagenorhynchus) obliquidens*, Rosser et al. 2022). Our recommendation may be useful for determining whether the social relationships found in these species are like alliances found in *Tursiops*.

## Data availability statement

The data that support the findings of this study are included in Table 1 and supplemental material.

## Acknowledgment

We sincerely thank Richard Connor for his insightful and constructive comments on our previous manuscript. We appreciate Takuya Aoki for drawing the illustrations to use in the figures. We would like to thank Editage (www.editage.jp) for English language editing.

## References

Allen SJ, King SL, Krützen M, Brown AM (2017) Multi-modal sexual displays in Australian humpback dolphins. Scientific Reports 7: 13644.

Aureli F, Schaffner CM, Boesch C, Bearder SK, Call J, Chapman CA et al. (2008) Fission-fusion dynamics: new research frameworks. Current Anthropology 49 (4): 627–654.

Baker I, O’Brien J, McHugh K, Berrow S (2020). Fine-scale sociality reveals female–male affiliations and absence of male alliances in bottlenose dolphins (*Tursiops truncatus*) in the Shannon Estuary, Ireland. Marine Mammal Science 36 (1): 66–88.

Bissonnette A, Franz M, Schülke O, Ostner J (2014) Socioecology, but not cognition, predicts male coalitions across primates. Behavioral Ecology 25 (4): 794–801.

Bissonnette A, Perry S, Barrett L, Mitani JC, Flinn M, Gavrilets S, de Waal FB (2015). Coalitions in theory and reality: a review of pertinent variables and processes. Behaviour 152 (1): 1–56.

Bizzozzero MR, Allen S.J, Gerber L, Wild S, King SL, Connor RC, Friedman WR, Wittwer S, Krützen M (2019). Tool use and social homophily among male bottlenose dolphins. Proceedings of the Royal Society B 286 (1904): 20190898.

Bouveroux TH, Mallefet J (2010). Social structure of bottlenose dolphins, *Tursiops truncatus*, in Panama City, Florida. Journal of the Marine Biological Association of the United Kingdom 90 (8): 1685–1692.

Brightwell K, Titcomb EM, Mazzoil M, Gibson Q (2020) Common bottlenose dolphin (*Tursiops truncatus*) social structure and distribution changes following the 2008 Unusual Mortality Event in the Indian River Lagoon, Florida. Marine Mammal Science 36 (4): 1271–1290.

Brightwell K, Gibson Q (2023) Inter-and intrapopulation variation in bottlenose dolphin mating strategies. In: Würsig B, Orbach DN (eds.) Sex in cetaceans: Morphology, behavior, and the evolution of sexual strategies, 251–278. Springer International Publishing, Cham, Switzerland.

Brightwell KK, Krzyszczyk EB, Gibson QA (2025) Dynamics of multilevel alliances in St. Johns River, Florida, Tamanend’s bottlenose dolphins (*Tursiops erebennus*) with respect to an epizootic unusual mortality event. Marine Mammal Science 41 (1), e13165.

Brusa JL, Young RF, Swanson T (2016) Abundance, ranging patterns, and social behavior of bottlenose dolphins (*Tursiops truncatus*) in an estuarine terminus. Aquatic Mammals 42 (1): 109–121.

Cairns SJ, Schwager SJ (1987) A comparison of association indices. Animal Behaviour 35 (5): 1454–1469.

Chabanne DB, Krützen M, Finn H, Allen SJ (2022) Evidence of male alliance formation in a small dolphin community. Mammalian Biology 102 (4): 1285–1298.

Chakrabarti S, Jhala YV (2017) Selfish partners: resource partitioning in male coalitions of Asiatic lions. Behavioral Ecology 28 (6): 1532–1539.

Chereskin E, Connor RC, Friedman WR, Jensen FH, Allen SJ, Sørensen PM, Krützen M and King SL (2022). Allied male dolphins use vocal exchanges to “bond at a distance”. Current Biology 32 (7): 1657–1663.

Chereskin E, Allen SJ, Connor RC, Krützen M, and King SL (2024). In pop pursuit: social bond strength predicts vocal synchrony during cooperative mate guarding in bottlenose dolphins. Philosophical Transactions B 379 (1905): 20230194.

Chilvers BL, Corkeron PJ (2001) Trawling and bottlenose dolphins’ social structure. Proceedings of the Royal Society B: Biological Sciences 268 (1479): 1901–1905.

Cioffi WR, Quick NJ, Foley HJ, Waples DM, Swaim ZT, Shearer JM et al. (2021) Adult male Cuvier’s beaked whales (*Ziphius cavirostris*) engage in prolonged bouts of synchronous diving. Marine Mammal Science 37 (3): 1085–1100.

Clutton-Brock T. (2016). Mammal Societies. Wiley, Oxford, UK.

Committee on Taxonomy (2023) List of marine mammal species and subspecies. Society for Marine Mammalogy. URL www.marinemammalscience.org. last consulted on 2025/01/08.

Connor RC, Smolker RA, Richards AF (1992a) Dolphin alliances and coalitions. In: Harcourt AH, de Waal F (eds.) Coalitions and alliances in humans and other animals, 415–443. Oxford University Press, Oxford, UK.

Connor RC, Smolker RA, Richards AF (1992b) Two levels of alliance formation among male bottlenose dolphins (*Tursiops* sp.). Proceedings of the National Academy of Sciences 89 (3): 987–990.

Connor RC, Richards AF, Smolker RA, Mann J (1996) Patterns of female attractiveness in Indian Ocean bottlenose dolphins. Behaviour 133 (1-2): 37–69.

Connor RC, Heithaus MR, Barre LM (1999) Superalliance of bottlenose dolphins. Nature 397 (6720): 571–572.

Connor RC, Wells RS, MannJ, Read AJ (2000) The bottlenose dolphin: social relationships in a fission-fusion society. In: Mann J, Conner RC, Tyack PL, Whitehead H (eds.) Cetacean societies: field studies of dolphins and whales, 91–126. University of Chicago Press, Chicago, USA.

Connor RC, Heithaus MR, Barre LM (2001) Complex social structure, alliance stability and mating access in a bottlenose dolphin ‘super-alliance’. Proceedings of the Royal Society of London. Series B: Biological Sciences 268 (1464): 263–267.

Connor RC, Smolker R, Bejder L (2006) Synchrony, social behaviour and alliance affiliation in Indian Ocean bottlenose dolphins, *Tursiops aduncus*. Animal Behaviour 72 (6): 1371–1378.

Connor RC, Mann J (2006) Social cognition in the wild: Machiavellian dolphins?. In: S. Hurley, M. Nudds (eds) Rational animals?, 329–367. Oxford University Press, Oxford, UK.

Connor RC (2007) Dolphin social intelligence: complex alliance relationships in bottlenose dolphins and a consideration of selective environments for extreme brain size evolution in mammals. Philosophical Transactions of the Royal Society B: Biological Sciences 362 (1480): 587–602.

Connor RC, Watson-Capps JJ, Sherwin WB, Krützen M (2011) A new level of complexity in the male alliance networks of Indian Ocean bottlenose dolphins (*Tursiops* sp.). Biology Letters 7 (4): 623–626.

Connor RC, Krützen M (2015) Male dolphin alliances in Shark Bay: changing perspectives in a 30-year study. Animal Behaviour 103 (1): 223–235.

Connor RC, Cioffi WR, Randić S, Allen SJ, Watson-Capps J, and Krützen M (2017). Male alliance behaviour and mating access varies with habitat in a dolphin social network. Scientific Reports 7: 46354.

Connor RC, Krützen M, Allen SJ, King SL (2022) Strategic intergroup alliances increase access to a contested resource in male bottlenose dolphins. Proceedings of National Academy of Sciences 119 (36): e2121723119.

Cords M, Thompson NA. Friendships, coalitions, and alliances. In: Call J, Burghardt GM, Pepperberg IM, Snowdon CT, Zentall T (eds.) APA handbook of comparative psychology: Basic concepts, methods, neural substrate, and behavior, 899-913. American Psychological Association, Washington DC, USA.

Croft P, James R, Krause J (2008) Exploring animal social networks. Princeton University Press, Princeton, USA

Danaher-Garcia N, Connor R, Fay G, Melillo-Sweeting K, Dudzinski M (2022) Using social network analysis to confirm the ‘gambit of the group’ hypothesis for a small cetacean. Behavioural Processes 200 (1): 104694.

Diaz-Aguirre F, Parra GJ, Passadore C, Möller L (2018) Kinship influences social bonds among male southern Australian bottlenose dolphins (*Tursiops* cf*. australis*). Behavioral Ecology and Sociobiology 72 (12): 190.

Duffield D, Wells R (2023) Paternity patterns in a long-term resident bottlenose dolphin community. Frontiers in Marine Science 10: 1076715.

Elliser CR, Herzing DL (2011) Replacement dolphins? Social restructuring of a resident pod of Atlantic bottlenose dolphins, *Tursiops truncatus*, after two major hurricanes. Marine Mammal Science 27 (1): 39–59.

Elliser CR, Herzing DL (2014) Long-term social structure of a resident community of Atlantic spotted dolphins, *Stenella frontalis*, in the Bahamas 1991–2002. Marine Mammal Science 30 (1): 308–328.

Ermak J, Brightwell K, Gibson Q (2017) Multi-level dolphin alliances in northeastern Florida offer comparative insight into pressures shaping alliance formation. Journal of Mammalogy 98 (4): 1096–1104.

Farine DR, Whitehead H (2015) Constructing, conducting and interpreting animal social network analysis. Journal of Animal Ecology 84 (5): 1144–1163.

Feldblum JT, Krupenye C, Bray J, Pusey AE, Gilby IC (2021) Social bonds provide multiple pathways to reproductive success in wild male chimpanzees. iScience 24 (8): 102864.

Félix F (1997) Organization and social structure of the coastal bottlenose dolphin *Tursiops truncatus* in the Gulf de Guayaquil, Ecuador. Aquatic Mammals 23 (1): 1–16.

Fox SA, Muller MN, González NT, Enigk DK, Machanda ZP, Otali E, Wrangham R, Thompson ME (2023). Weak, but not strong, ties support coalition formation among wild female chimpanzees. Philosophical Transactions of the Royal Society B 378 (1868): 20210427.

Frau S, Ronchetti F, Perretti F, Addis A, Ceccherelli G, La Manna G (2021) The influence of fish farm activity on the social structure of the common bottlenose dolphin in Sardinia (Italy). PeerJ 9: e10960.

Friedman WR, Krützen M, King SL, Allen SJ, Gerber L, Wittwer S, Connor RC (2023) Inter-group alliance dynamics in Indo-Pacific bottlenose dolphins (*Tursiops aduncus*). Animal Cognition 26 (5): 1601–1612.

Genov T, Centrih T, Kotnjek P, Hace A (2019) Behavioural and temporal partitioning of dolphin social groups in the northern Adriatic Sea. Marine Biology 166 (1): 11.

Gerber L, Connor RC, King SL, Allen SJ, Wittwer S, Bizzozzero MR et al. (2020) Affiliation history and age similarity predict alliance formation in adult male bottlenose dolphins. Behavioral Ecology 31 (2): 361–370.

Gerber L, Wittwer S, Allen SJ, Holmes KG, King SL, Sherwin WB, Wild S, Willems EP, Connor RC, Krützen M (2021) Cooperative partner choice in multi-level male dolphin alliances. Scientific Reports 11: 6901.

Gerber L, Connor RC, Allen SJ, Horlacher K, King SL, Sherwin WB, Willems EP, Wittwer S, Krützen M (2022) Social integration influences fitness in allied male dolphins. Current Biology 32 (7): 1664–1669.

Gilby IC, Brent LJ, Wroblewski EE, Rudicell RS, Hahn BH, Goodall J, Pusey AE (2013) Fitness benefits of coalitionary aggression in male chimpanzees. Behavioral Ecology and Sociobiology 67 (3): 373–381.

Gowans S (2019) Grouping behaviors of dolphins and other toothed Whales. In: Würsig B (ed.) Ethology and Behavioral Ecology of Odontocetes, 3–24. Springer, Cham, Switzerland.

Harcourt AH, de Waal F (1992). Coalitions and alliances in humans and other animals. Oxford University Press, Oxford, UK

Hartman KL, Visser F, Hendriks AJ (2008) Social structure of Risso’s dolphins (*Grampus griseus*) at the Azores: a stratified community based on highly associated social units. Canadian Journal of Zoology 86 (4): 294–306.

Hartman K, Van der Harst P, Vilela R (2020) Continuous focal group follows operated by a drone enable analysis of the relation between sociality and position in a group of male Risso’s dolphins (*Grampus griseus*). Frontiers in Marine Science 7: 283.

Hartman KL, Chen I, van der Harst PA, Moura AE, Jahnke M, Pilot M, Vilela PR, Hoelzel AR (2023) Kinship study reveals stable non-kin-based associations in a medium-sized delphinid. Behavioral Ecology and Sociobiology 77 (12): 137.

Herzing DL (1996) Vocalizations and associated underwater behavior of free-ranging Atlantic spotted dolphins, *Stenella frontalis* and bottlenose dolphins, *Tursiops truncatus*. Aquatic Mammals 22 (2): 61–80.

Herzing DL, Johnson CM (1997) Interspecific interactions between Atlantic Spotted dolphins (*Stenella frontalis*) and Bottlenose dolphins (*Tursiops truncatus*) in the Bahamas, 1985–1995. Aquatic Mammals 23 (2): 85–99.

Herzing DL, Moewe K, Brunnick BJ (2003) Interspecies interactions between Atlantic spotted dolphins, *Stenella frontalis* and bottlenose dolphins, *Tursiops truncatus*, on Great Bahama Bank, Bahamas. Aquatic Mammals 29 (3): 335–341.

Herzing DL, Elliser CR (2013) Directionality of sexual activities during mixed-species encounters between Atlantic spotted dolphins (*Stenella frontalis*) and bottlenose dolphins (*Tursiops truncatus*). International Journal of Comparative Psychology 26 (2): 124–134.

Hill-Cousins S, Chereskin E, Allen SJ, Connor RC, Krützen M, Papageorgiou D, King SL (2025). Allied male dolphins use synchronous displays to strengthen social bonds in a cooperative context. Movement Ecology 13 (1): 84.

Hoppitt WJ, and Farine DR (2018). Association indices for quantifying social relationships: how to deal with missing observations of individuals or groups. Animal Behaviour 136 (1): 227–238.

Janik VM (2000). Source levels and the estimated active space of bottlenose dolphin (Tursiops truncatus) whistles in the Moray Firth, Scotland. Journal of Comparative Physiology A 186 (7): 673–680.

Jennings DJ, Carlin CM, Hayden TJ, Gammell MP (2011) Third-party intervention behaviour during fallow deer fights: the role of dominance, age, fighting and body size. Animal Behaviour 81 (6): 1217–1222.

Johnson CM, Norris KS (1994) Social behavior. In: Norris KS, Würsig B, Wells RS, Würsig M (eds.) The Hawaiian Spinner Dolphin, 243–286. University of California Press, Berkeley, CA.

Kawazoe T (2021) Male–male social bonds predict tolerance but not coalition formation in wild Japanese macaques. Primates 62 (1): 91–101.

Kent EE, Mazzoil M, McCulloch SD, Defran RH (2008) Group characteristics and social affiliation patterns of bottlenose dolphins (*Tursiops truncatus*) in the Indian River Lagoon, Florida. Florida Scientist 71 (2): 149–168.

King SL, Friedman WR, Allen SJ, Gerber L, Jensen FH, Wittwer S, Connor RC. Krützen M (2018) Bottlenose dolphins retain individual vocal labels in multi-level alliances. Current Biology 28 (12): 1993–1999.

King SL, Connor RC, Krützen M, Allen SJ (2021) Cooperation-based concept formation in male bottlenose dolphins. Nature Communications 12 (1): 2373.

King SL and Jensen FH (2023). Rise of the machines: Integrating technology with playback experiments to study cetacean social cognition in the wild. Methods in Ecology and Evolution 14 (8): 1873–1886.

Kobayashi H, Whitehead H, Amano, M (2020) Long-term associations among male sperm whales (*Physeter macrocephalus*). PLoS One 15: e0262066.

Krützen M, Barré LM, Connor RC, Mann J, Sherwin WB (2004) ‘O father: where art thou?’ - Paternity assessment in an open fission–fusion society of wild bottlenose dolphins (*Tursiops* sp.) in Shark Bay, Western Australia. Molecular Ecology 13 (7): 1975–1990.

Kutsukake N, Hasegawa T (2005) Dominance turnover between an alpha and a beta male and dynamics of social relationships in Japanese macaques. International Journal of Primatology 26 (4): 775–800.

Lukas D, Clutton-Brock T (2018) Social complexity and kinship in animal societies. Ecology Letters 21 (8): 1129–1134.

Lusseau D, Schneider K, Boisseau OJ, Haase P, Slooten E, Dawson SM (2003) The bottlenose dolphin community of Doubtful Sound features a large proportion of long-lasting associations - Can geographic isolation explain this unique trait? Behavioral Ecology and Sociobiology 54 (4): 396–405.

Lusseau D (2007) Why are male social relationships complex in the Doubtful Sound bottlenose dolphin population? PLoS One 4: e348.

Mammal Diversity Database (2023) Mammal Diversity Database (Version 1.12.1) [Data set]. Zenodo. URL 10.5281/zenodo.10595931

Mann J (1999) Behavioral sampling methods for cetaceans: a review and critique. Marine Mammal Science 15 (1): 102–122.

Maze-Foley K, Würsig B (2002) Patterns of social affiliation and group composition for bottlenose dolphins (*Tursiops truncatus*) in San Luis Pass, Texas. Gulf of Mexico Science 20 (2): 122–134.

Möller LM, Beheregaray LB, Harcourt RG, Krützen M (2001) Alliance membership and kinship in wild male bottlenose dolphins (*Tursiops aduncus*) of southeastern Australia. Proceedings of the Royal Society B 268 (1479): 1941–1947.

Möller LM (2012) Sociogenetic structure, kin associations and bonding in delphinids. Molecular Ecology 21 (3): 745–764.

Moreno K, Acevedo-Gutiérrez A (2016) The social structure of Golfo Dulce bottlenose dolphins (*Tursiops truncatus*) and the influence of behavioural state. Royal Society Open Science 3 (8): 160010.

Morteo E, Rocha-Olivares A, Abarca-Arenas LG (2014) Sexual segregation in coastal bottlenose dolphins (*Tursiops truncatus*) in the south-western Gulf of Mexico. Aquatic Mammals 40 (4): 375–385.

Nishita M, Shirakihara M, Iwasa N, Amano M (2017) Alliance formation of Indo-Pacific bottlenose dolphins (*Tursiops aduncus*) off Amakusa, western Kyushu, Japan. Mammal Study 42 (3): 125–130.

O’Dea RE, Lagisz M, Jennions MD, Koricheva J, Noble DW, Parker TH, Gurevitch J, Page MJ, Stewart G, Moher D, Nakagawa, S (2021). Preferred reporting items for systematic reviews and meta-analyses in ecology and evolutionary biology: a PRISMA extension. Biological Reviews 96 (5): 1695–1722.

Olson LE, Blumstein DT (2009) A trait-based approach to understand the evolution of complex coalitions in male mammals. Behavioral Ecology 20 (3): 624–632.

Owen EC, Wells RS, Hofmann S (2002) Ranging and association patterns of paired and unpaired adult male Atlantic bottlenose dolphins, *Tursiops truncatus*, in Sarasota, Florida, provide no evidence for alternative male strategies. Canadian Journal of Zoology 80 (12): 2072–2089.

Parsons KM, Durban JW, Claridge DE (2003a) Male-male aggression renders bottlenose dolphin (*Tursiops truncatus*) unconscious. Aquatic Mammals 29 (3): 360–362.

Parsons KM., Durban JW, Claridge DE, Balcomb KC, Noble LR, Thompson PM (2003b) Kinship as a basis for alliance formation between male bottlenose dolphins, *Tursiops truncatus*, in the Bahamas. Animal Behaviour 66 (1): 185–194.

Pryor K, Kang-Shallenberger I (1991) Social structure in spotted dolphins (*Stenella attenuata*) in the tuna purse seine fishery in the eastern Tropical Pacific. In: Pryor K, Norris KS (eds.) Dolphin societies: discoveries and puzzles, 161–196. University of California Press, Berkeley, CA.

Quintana-Rizzo E, Mann DA, Wells RS (2006). Estimated communication range of social sounds used by bottlenose dolphins (*Tursiops truncatus*). The Journal of the Acoustical Society of America 120 (3): 1671–1683.

Rako-Gospić N, Radulović M, Vučur T, Pleslić G, Holcer D, Mackelworth P (2017) Factor associated variations in the home range of a resident Adriatic common bottlenose dolphin population. Marine Pollution Bulletin 124 (1): 234–244.

Randić S, Connor RC, Sherwin WB, Krützen M (2012) A novel mammalian social structure in Indo-Pacific bottlenose dolphins (*Tursiops* sp.): complex male alliances in an open social network. Proceedings of the Royal Society B: Biological Sciences 279 (1740): 3083–3090.

Rogers CA, Brunnick BJ, Herzing DL, Baldwin JD (2004) The social structure of bottlenose dolphins, *Tursiops truncatus*, in the Bahamas. Marine Mammal Science 20 (4): 688–708.

Rosser LR, Morisaka T, Mitani Y, Igarashi T. (2022). Calf-directed aggression as a possible infanticide attempt in Pacific white-sided dolphins (*Lagenorhynchus obliquidens*). Aquatic Mammals 48 (3): 273–286.

Samuels A, Tyack P (2000) Flukeprints: A history of studying cetacean societies. In: Mann J, Conner RC, Tyack PL, Whitehead H (eds.) Cetacean societies: field studies of dolphins and whales, 9–44. University of Chicago Press, Chicago, USA.

Scott EM, Mann J, Watson-Capps JJ, Sargeant BL, Connor RC (2005) Aggression in bottlenose dolphins: evidence for sexual coercion, male-male competition, and female tolerance through analysis of tooth-rake marks and behaviour. Behaviour 142 (1): 21–44.

Smith JE, Fichtel C, Holmes RK, Kappeler PM, van Vugt M, Jaeggi AV (2022) Sex bias in intergroup conflict and collective movements among social mammals: male warriors and female guides. Philosophical Transactions of the Royal Society B 377 (1851): 20210142.

Smith JE, Jaeggi AV, Holmes RK, Silk JB (2023) Sex differences in cooperative coalitions: a mammalian perspective. Philosophical Transactions of the Royal Society B 378 (1868): 20210426.

Smith JE, Van Horn RC, Powning KS, Cole AR, Graham KE, Memenis SK, Holekamp KE (2010) Evolutionary forces favoring intragroup coalitions among spotted hyenas and other animals. Behavioral Ecology 21 (2): 284–303.

Smolker RA, Richards AF, Connor RC, Pepper JW (1992) Sex differences in patterns of association among Indian Ocean bottlenose dolphins. Behaviour 123 (1-2): 38–69.

Sørensen PM, Connor RC, Allen SJ, Krützen M, Lebrec U, Jensen FH, King SL (2024). Communication range predicts dolphin alliance size in a cooperative mating system. Current Biology 34 (20): 4774–4780.

Sterck EH, Watts DP, Van Schaik CP (1997) The evolution of female social relationships in nonhuman primates. Behavioral Ecology and Sociobiology 41 (5): 291–309.

Strauss ED, Holekamp KE (2019) Social alliances improve rank and fitness in convention-based societies. Proceedings of the National Academy of Sciences 116 (18): 8919–8924.

Syme J, Kiszka JJ, Parra GJ (2022). How to define a dolphin “group”? Need for consistency and justification based on objective criteria. Ecology and Evolution 12 (11): e9513.

Tokuyama N, Furuichi T (2016) Do friends help each other? Patterns of female coalition formation in wild bonobos at Wamba. Animal Behaviour 119 (1): 27–35.

Toyoda A, Maruhashi T, Kawamoto Y, Matsudaira K, Matsuda I, Malaivijitnond S (2022) Mating and reproductive success in free-ranging stump-tailed macaques: effectiveness of male–male coalition formation as a reproductive strategy. Frontiers in Ecology and Evolution 10: 802012.

van Schaik (1989) The ecology of social relationships amongst female primates. In: Standen V, Foley RA (eds.) Comparative Socioecology: The behavioural ecology of humans and other mammals, 195–218. Blackwell, Oxford, UK.

Vermeulen E (2018) Association patterns of bottlenose dolphins (*Tursiops truncatus*) in Bahía San Antonio, Argentina. Marine Mammal Science 34 (3): 687–700.

Wallen MM, Patterson EM, Krzyszczyk E, Mann J (2016) The ecological costs to females in a system with allied sexual coercion. Animal Behaviour 115 (1): 227–236.

Weiss MN, Franks DW, Croft DP, Whitehead H (2019) Measuring the complexity of social associations using mixture models. Behavioral Ecology and Sociobiology 73 (1): 8.

Weiss MN, Franks DW, Giles DA, Youngstrom S, Wasser SK, Balcomb KC et al. (2021) Age and sex influence social interactions, but not associations, within a killer whale pod. Proceedings of the Royal Society B 288 (1953): 20210617.

Wells RS, Scott MD, Irvine AB (1987) The social structure of free-ranging bottlenose dolphins. In: Genoways HH (ed.) Current Mammalogy, 247–305. Springer, New York, USA.

Wells RS (2014) Social structure and life history of bottlenose dolphins near Sarasota Bay, Florida: insights from four decades and five generations. In: Yamagiwa J, Karczmarski L (eds.) Primates and Cetaceans: Field Research and Conservation of Complex Mammalian Societies, 149–172. Springer, New York, Japan.

Whitehead H, Connor RC (2005) Alliances I. How large should alliances be? Animal Behaviour 69 (1): 117–126.

Whitehead H (2008). Analyzing animal societies: quantitative method for vertebrate social analysis. University of Chicago Press, Chicago, USA.

Whitehead H and James R (2015) Generalized affiliation indices extract affiliations from social network data. Methods in Ecology and Evolution 6 (7): 836–844.

Wilson DE, Reeder DM (2005). Mammal species of the world. A taxonomic and geographic reference, 3rd ed. Johns Hopkins University Press, Baltimore, USA.

Wiszniewski J, Brown C, Möller LM (2012a) Complex patterns of male alliance formation in a dolphin social network. Journal of Mammalogy 93 (1): 239–250.

Wiszniewski J, Corrigan S, Beheregaray LB, Möller LM (2012b) Male reproductive success increases with alliance size in Indo-Pacific bottlenose dolphins (*Tursiops aduncus*). Journal of Animal Ecology 81 (2): 423–431.

