## supplemental data for "Methodological inconsistencies hinder comparative studies of alliance in the genus *Tursiops*"

Connor RC, Smolker RA. (1996). 'Pop 'goes the dolphin: A vocalization male bottlenose dolphins produce during consortships. *Behaviour*: 133 (9-10): 643-662.

Connor RC, Heithaus MR, Barre LM (1999) Superalliance of bottlenose dolphins. *Nature* 397: 571-572.

King SL, Friedman WR, Allen SJ, Gerber L, Jensen FH, Wittwer S, Connor RC. Krützen M (2018) Bottlenose dolphins retain individual vocal labels in multi-level alliances. *Current Biology* 28 (12): 1993-1999.

King SL, Allen SJ, Krützen M, Connor RC (2019). Vocal behaviour of allied male dolphins during cooperative mate guarding. *Animal cognition* 22 (6): 991-1000.

Vermeulen E (2018) Association patterns of bottlenose dolphins (*Tursiops truncatus*) in Bahía San Antonio, Argentina. *Marine Mammal Science* 34 (3): 687-700.

Vollmer NL, Hayek LAC, Heithaus MR, Connor RC (2015). Further evidence of a context-specific agonistic signal in bottlenose dolphins: the influence of consortships and group size on the pop vocalization. *Behaviour*: 152 (14): 1979-2000.

Wells RS, Scott MD, Irvine AB (1987) The social structure of free-ranging bottlenose dolphins. In: Genoways HH (ed.) Current Mammalogy, 247-305. Springer, New York, USA.

Wells RS (2014) Social structure and life history of bottlenose dolphins near Sarasota Bay, Florida: insights from four decades and five generations. In: Yamagiwa J, Karczmarski L (eds.) Primates and Cetaceans: Field Research and Conservation of Complex Mammalian Societies, 149-172. Springer, New York, Japan.

Wiszniewski J, Brown C, Möller LM (2012) Complex patterns of male alliance formation in a dolphin social network. *Journal of Mammalogy* 93 (1): 239-250.

Table S1: Definitions of association, group and association index for each study.

| Alliance categories | Regions & species | Definition of association | Definition of group | Association index |
|---|---|---|---|---|
| Alliance formation | Shark Bay (*T. aduncus*) | Group membership [1-6,8]<br>Consortship†1 [13,14] | Proximity* [3-5,7,10,11]<br>Combination (proximity* & behavioral state†2) [1,2,8,9] | SRI [1-7]<br>HWI [8-12] |
| | Port Stephens (*T. aduncus*) | Group membership [15,16] | Combination (proximity+ & behavioral state†3) [15,16] | HWI [15,16] |
| | Amakusa – Shimoshima (*T. aduncus*) | Proximity (Within 3 body length) [17] | - | HWI [17] |
| | Swan Canning Riverpark (*T. aduncus*) | Group membership [18] | Combination (proximity* & behavioral state†3) [18] | SRI [18] |
| | Sarasota Bay (*T. truncatus*) | Group membership [19,20] | Proximity+ [19]<br>Combination (proximity+ & behavioral state†3) [20] | HWI [19,20] |
| | St. Johns River (*T. truncatus*) | Group membership [21] | Proximity* [21,22] | HWI [21,22] |
| | Abaco Island (*T. truncatus*) | Group membership [23] | No info | HWI [23] |

| | | | | |
|---|---|---|---|---|
| | Indian River Lagoon (*T. truncatus*) | No info | Combination (proximity+ & behavioral state†3) [24,25] | HWI [24,25] |
| | Cres-Lošinj archipelago (*T. truncatus*) | No info | No info | HWI [26] |
| | Little Bahama Bank (*T. truncatus*) | Group membership [27] | Behavioral state# [27,28] | HWI [27,28] |
| | Gulf of Guayaquil (*T. truncatus*) | No info | No info | HWI [29] |
| | Panama City (*T. truncatus*) | Group membership [30] | Combination (proximity+ & behavioral state†3) [30] | HWI [30] |
| Putative alliance formation | Moreton Bay (*T. aduncus*) | Group membership [31] | Proximity* [31] | HWI [31] |
| | Golfo Dulce (*T. truncatus*) | Group membership [32] | Proximity* [32] | HWI [32] |
| | San Luis Pass (*T. truncatus*) | Group membership [33] | Proximity* [33] | HWI [33] |

| | | | | |
|---|---|---|---|---|
| | Doubtful Sound (*T. truncatus*) | Group membership [35] | Combination (proximity & behavioral state†3) [34,35] | HWI [34-35] |
| | Alvarado Lagoon (*T. truncatus*) | Group membership [36] | Behavioral state# [36] | HWI [36] |
| | Coffin Bay (*Tursiops.* sp) | No info | Combination (proximity+ & behavioral state†3) [37] | GAI [37]<br>HWI [37] |
| Non-alliance formation | Shannon Estuary (*T. truncatus*) | Group membership [38] | Combination (proximity+ & behavioral state†3 & sighting together) [38] | HWI [38] |
| | Gulf of Trieste (*T. truncatus*) | Group membership [39] | Combination (proximity+ & behavioral state†3) [39] | HWI [39] |
| | Bahía San Antonio (*T. truncatus*) | Group membership [40] | Combination (proximity+ & behavioral state†3) [40] | HWI [40] |
| | North Inlet-Winyah Bay (*T. truncatus*) | Group membership [41] | Combination (proximity$ & behavioral state†3) [41] | HWI [41] |
| | Sardinia (*T. truncatus*) | Group membership [42] | Combination (proximity & behavioral state†3 & sighting together) [42] | HWI [42] |

Reference numbers correspond to the following; [1] Smolker et al. (1992), [2] Connor et al. (1992), [3] Connor et al. (2011), [4] Randić et al. (2012), [5] Bizzozzero et al. (2019), [6] Gerber et al. (2020), [7] Friedman et al. (2023), [8] King et al.

(2018), [9] Connor et al. (2022), [10] King et al. (2021), [11] Gerber et al. (2022), [12]Gerber et al. (2021), [13] Connor et al. (1999), [14] Connor et al. (2001), [15] Möller et al. (2001), [16] Wiszniewski et al. (2012), [17] Nishita et al. (2017), [18 ] Chabanne et al. (2022), [19] Wells et al. (1987), [20] Owen et al. (2002) , [21] Ermak et al. (2017), [22] Brightwell et al. (2024), [23] Parsons et al. (2003), [24] Kent et al. (2008), [25] Brightwell et al. (2020), [26] Rako-Gospić et al. (2017), [27] Rogers et al. (2004), [28] Elliser and Herzing (2011), [29] Félix (1997), [30] Bouveroux and Mallefet (2010), [31] Chilvers and Corkeron (2001), [32] Moreno and Acevedo-Gutiérrez (2016), [33] Maze-Foley and Würsig (2002), [34] Lusseau et al. (2003), [35] Lusseau (2007), [36] Morteo et al. (2014), [37] Diaz-Aguirre et al. (2018), [38] Barker et al. (2020), [39] Genov et al. (2019), [40] Vermeulen (2018), [41] Brusa et al. (2016), [42] Frau et al. (2021).

*: 10m-chain rule (see Smolker et al. 1992). +: 100m radius rule (see Wells et al. 1987). #: behavioral coordination rule (see Shane 1990). $: proximity threshold set as 40m. †1: specific state data are included in the analysis (especially, consortship state data are only used). †2: specific state data are included in the analysis (especially, foraging state data are not used). †3: same behavioral state such as direction of swimming or same behavioral state such as traveling and resting.

Table S2. Behaviors observed among males with stable associations. Type indicates interaction directionality: “M to F” denotes male-to-female interactions, and “M to M” denotes male-to-male interactions.

| Regions & Species | Type | Description |
| --- | --- | --- |
| Shark Bay (*T. aduncus*) | M to F<br>M to M | Because first-order alliances differ from second- and third-order alliances in their primary functions (Connor and Krützen 2015; Connor et al. 2022), we here focus on first-order alliances. The primary function of first-order alliances is the maintenance of consortship with females. Consortship is recorded when the behavioral elements described below are observed:<br>1. Capture: “A 'capture' was recorded if the consortship was initiated when the males traveled rapidly toward the female (typically leaping or porpoising), then porpoised around her or engaged with her in intense social behaviour that included body-body contact.” (Connor et al. 1996, Results)<br>2. Bolt: “Recording a 'bolt' required evidence that the female accelerated first, which usually included observation of the female accelerating or the trail of disturbed surface water that was produced by her rapid swimming. Bolts could also be inferred if we saw the males accelerate and the female already had gained a significant lead in the chase indicating that she had accelerated before the male” (Connor et al. 1996, Results. See also Connor and Krützen 2015, supplemental materials).<br>3. Pop: “We only scored 'pops' produced in air at the surface where we could identify the vocalizing individual, or at least narrow it down to one of the males.... Pops produced in air were easy to localize and movement of the blowhole and water sputtering from the blowhole often provided a visible |

manifestation of pops." (Connor et al. 1996, Results. See also See also Connor and Krützen 2015, supplemental materials; Vollmer et al. 2015; King et al. 2019). "Pops are narrow-band, low frequency pulses with peak energy between 300 and 3000 Hz and are typically produced in trains of 3-30 pops at rates of 6-12 pops/s." (Connor and Smolker 1996, Abstract)

4. Aggression: "…physical threats or aggression by the males toward the female. The aggressive behaviours we recorded were charging, 'head-jerks' (rapid vertical or lateral movements of the head), or hitting with the tail." (Connor et al. 1996, Results). "Documented aggression requires observation of the males threating the female with 'head-jerks, charging at the female or hitting her..." (Connor and Krützen 2015, supplemental materials).

5. Theft: "an attempted theft of the female by other alliances" (Connor and Krützen 2015, supplemental materials. But see Connor et al. 1996, Results)

6. Long-lasting association: "an association between the alliances and the female for at least 1 h during follows or during multiple surveys spanning 1 h" (Connor and Krützen 2015, supplemental materials).

The term consortship, in a general sense, refers simply to an association between a male and a female. However, long-term studies have demonstrated that such male-female associations are actively maintained by males, who play a dominant role in sustaining these interactions. Accordingly, in Shark Bay, the term consortship is used to refer specifically to coerced consortship (also termed herding).

| | | |
|---|---|---|
| Shark Bay (*T. aduncus*) | M to M | Here we focus on second- and third-order alliances. The primary function of these higher-order alliances is cooperation against rival groups over access to females, including behaviours such as female guarding and theft (Connor and Krützen 2015, Connor et al. 2022). Detailed observations of second-order alliances are reported in Connor et al. (1992a,b), and detailed descriptions of third-order alliances are provided in Connor et al. (2011). |
| Port Stephens (*T. aduncus*) | M to F | “Individuals were considered herding if we observed a \`captured female’, a \`capture attempt’ or an \`escape attempt’ (Connor *et al*. 1992*b*). Males with a captured female usually travelled tightly positioned on either side of her and/or behind her. Capture attempts involved males chasing a female from the sides and behind her or rushing up and around her. An escape attempt was characterized by a captured female bolting away from the males” (Möller et al. 2001, Methods). |
| Amakusa - Shimoshima (*T. aduncus*) | M to F | “Of all photographs in which the males of the possible alliance members were identified, we confirmed 30 cases (one case refers to a series of events in a single sampling session) in which nine of the above-mentioned ten male pairs surrounded a female dolphin (Table 3, Fig. 2). In 17 of these 30 cases, females surrounded by male pairs were considered to be receptive at that time and eight females gave birth in the following year (Table 3).” (Nishita et al. 2017, Results). |

| | | |
|---|---|---|
| Swan Canning Riverpark (*T. aduncus*) | M to F | "A triad of allied SCR resident males herding a female that is resident to adjacent coastal waters in the estuary. The allied males travelled 'in formation' behind the female.", "Behavioural observations of the 'rooster strut' display (seen as a series of consecutive photos) performed by a SCR resident male in the presence of a female. As the male is at the surface, his head is arched above the surface and bobbed up and down while moving forward" (Chabanne et al. 2022, supplemental materials). |
| Sarasota Bay (*T. truncatus*) | M to F | "On several occasions, a male pair has been observed apparently separating an individual female from a school." (Wells et al. 1987, Discussion). It is noted that "Aggressive interactions between males and females in reproductive contexts appear to be much less common in Sarasota Bay than at other sites where male alliances have been observed……" (Wells 2014, Discussion). |
| St. Johns River (*T. truncatus*) | M to F | "…all 30 allied MAL-UNK have been documented in herding formation, surfacing synchronously while flanking a known female (Connor et al. 1996)." (Ermak et al. 2017, Results). "Consortships were identified when males consistently surfaced synchronously and swam abreast or flanked a known female on either side of it. Allied males were also often documented swimming tightly together on one side of the female rather than flanking during consortships. Consortships were also documented if a capture attempt of a female, escape attempt by the female, or physical aggression towards the female were observed (adapted from Connor et al., 1992, 1996; Möller et al., 2001; Watson-Capps, 2005)." (Brightwell et al. 2025, Methods). |

| | | |
|---|---|---|
| Abaco Island (*T. truncatus*) | M to M | "Putative male alliances were identified from repeated dolphin encounters and ad libitum observations (unpublished data). Behaviours that were definitive of alliances included 'herding' or attempted herding of females (with or without dependent calves), 'formation' swimming (Connor et al. 1992), synchronous or apparently coordinated behaviours, and coordinated aggressive behaviours directed at conspecifics without the alliance." (Parsons et al. 2003a, Method). "Herein, we describe a unique encounter involving a long-term male alliance competing with a 'solo' male that resulted in the temporary loss of consciousness of the lone male following repeated physical blows to his head region. This observation supports the increased fitness experienced by males in alliances and illustrates the potential severity of aggressive interactions among adult bottlenose dolphins." (Parsons et al. 2003b). |
| Little Bahama Bank (*T. truncatus*) | M to M[#] M to F[#] | "…the two male bottlenose dolphins then pursued and engaged another young adult male spotted dolphin. Similar sexual behavior occurred between these individuals. During this encounter head-to-head posturing, squawking, and other aggressive displays (by both species) were observed. The bottlenose dolphins were successful in mounting and copulating with the spotted male dolphins and reciprocal sexual behavior was not observed. Throughout the next four years, these types of interspecies encounters between males were observed dozens of times (Figs 7 and 8). Typically, interactions, which included penile intromission attempts or actual intromission, involved the young spotted dolphins taking a passive role, floating without resistance while the adult bottlenose dolphin |

| | | |
|---|---|---|
| | | actively arched and rubbed its body and genitals against the young spotted dolphin." (Herzing and Johnson 1997, Results). "…interspecific (vs intraspecific) interactions were observed across species in our study. In one case, male spotted dolphins joined male bottlenose dolphins pursuing a female bottlenose dolphin and, in another, one bottlenose dolphin joined a male spotted dolphin coalition pursuing a female spotted. It is interesting to note, however, that in both cases, the subsequent mating behavior was only intraspecies." (Herzing and Johnson 1997, Discussion). |
| Gulf of Guayaquil (*T. truncatus*) | M to M | "Twice, males #108 and #109 from community #2 were observed attacking other dolphins: chasing, head-on collision, back leaps to fall over the other dolphin and other obvious and violent movements not commonly observed." (Félix 1997, Results). |
| San Luis Pass (*T. truncatus*) | M to F | "In SLP, it has not been determined whether male pairs or trios herd females; however, we made observations on three different days that resembled descriptions of herding attempts in Shark Bay." (Maze-Foley and Würsig 2002, Discussion). |
| Doubtful Sound (*T. truncatus*) | M to M | "On five occasions I was able to identify all opponents of headbutting bouts involving three individuals (Table 1). All coalitions were formed of individuals from the same school. In two cases the two individuals that joined forces to fight another were from the same group but were not immediate associates (Gallatin and Jet against Jonah, Gallatin and SN90 against PL, Figures 1 and 2). In the three remaining instances the two allied individuals were pairs that were identified as |

| | | |
|---|---|---|
| | | spending the most time together (pairs with the highest association index for each individual involved in the pair).” (Lusseau 2007, Results). |
| Shannon Estuary (*T. truncatus*) | - | “Additionally, during observations of dolphin behavior, there were no groups of dolphins that appeared to behave in any ways typical of male alliances recorded in other study sites. For example, we did not observe chasing, bolting, displays, or aggression (e.g., charging or biting) that are often typical components of the behaviors exhibited by male alliances and the females they are herding during consortship behavior in Shark Bay (Baker et al., 2017; Connor and Krützen, 2015; Connor et al., 1992). Similarly, we never recorded any instances of dolphins in the Shannon Estuary producing popping vocalizations, nor did the dolphins exhibit “formation swimming” typical of that shown by male alliances during herding behavior (males traveling just behind and to either side of the herded female; Connor et al., 1992). This lack of behavioral evidence coupled with the reported results from data analysis support the idea that the Shannon Estuary bottlenose dolphins may not form male alliances.” (Barker et al. 2020, Discussion). |

#: inter-species copulation and competition were observed.
